# The methane-cycling microbiome in intact and degraded permafrost soils of the pan-Arctic

**DOI:** 10.1101/2024.12.16.628686

**Authors:** Haitao Wang, Erik Lindemann, Patrick Liebmann, Milan Varsadiya, Mette Marianne Svenning, Muhammad Waqas, Sebastian Petters, Andreas Richter, Georg Guggenberger, Jiri Barta, Tim Urich

## Abstract

The methane-cycling microbiomes in Arctic permafrost-affected soils play crucial roles in the production and consumption of this important greenhouse gas. However, little is known about the distributions of Arctic methanogens and methanotrophs across the regional scale and along the vertical soil profile, as well as their responses to the widespread permafrost thaw. Using a unique sample set from nine different locations across the pan-Arctic, we identified methanogen and methanotroph phylotypes in 729 datasets of 16S rRNA gene amplicons.

In 621 samples of intact permafrost soils across the pan-Arctic, only 22 methanogen and 26 methanotroph phylotypes were identified. Relative abundances of both functional groups varied significantly between sites and soil horizons. Only four methanogen phylotypes were detected at all locations, with the hydrogenotrophic *Methanobacterium lacus* dominating. Remarkably, the permafrost soil methane filter was almost exclusively comprised of a few phylotypes closely related to the obligate methanotrophic species *Methylobacter tundripaludum*.

In degraded permafrost sites in Alaska, *M. tundripaludum* also dominated the methanotroph microbiome in the wet site. However, in dry, water-drained former permafrost site, *Methylocapsa* phylotypes, closely related with the atmospheric methane oxidizing bacteria, were exclusively found and dominant, indicating a massive restructuring of the methanotroph guild that consequently resulted in functional changes from a soil methane filter to an atmospheric methane sink.

This study provides first insights into the identity and intricate spatial distribution of methanotrophs and methanogens in permafrost soils at a pan-Arctic scale and their responses to different water status after permafrost degradation. These findings point towards a few key microbes particularly relevant for future studies on Arctic CH_4_ dynamics in a warming climate and that under future dry conditions more atmospheric CH_4_ uptake in Arctic upland soils might happen.

## Introduction

Permafrost is primarily distributed across the Arctic region, underlying ∼25% of the land area in the Northern Hemisphere and hosting an estimated carbon stock of 1500-1700 Pg [1, 2]. Permafrost thaw due to climate change leads to the thickening of the active layer, which releases the frozen carbon and increases the availability and mobility of soil organic carbon (SOC) [3, 4]. This ongoing effect accelerates the potential of microbial organic matter decomposition and is thought to increase emissions of greenhouse gases, CH_4_ and CO_2_ [2, 5], which may result in a positive feedback effect into the climate system [1]. Depending on ice richness and soil drainage, permafrost thaw can result in water-saturated soils due to the abrupt collapse of ice wedges [6, 7], or drier soils due to better drainage and evapotranspiration [8, 9]. While drier and aerated soils may host a burst of CO_2_ via the microbial degradation of ancient carbon [1, 10], anoxic conditions in wetter soils may hamper this process and contribute to SOC accumulation [11]. However, anoxia in water-saturated soils favors anaerobic methanogenesis and the expansion of anoxic conditions after permafrost thaw may enhance the proportion of future CH_4_ emissions [12].

CH_4_ has ∼34 times higher warming potential compared to CO_2_ on a 100-year timescale [13]. Biogenic CH_4_ is primarily produced by methanogenic archaea carrying out anaerobic methanogenesis, despite some bacteria and eukaryotes producing CH_4_ using different mechanisms [14–16]. Anaerobic methanogenic archaea constitute the major archaeal communities in permafrost soils [17, 18]. Typical methanogens are a diverse archaeal group occurring within 8 validly described orders, *Methanobacteriales*, *Methanococcales*, *Methanopyrales*, *Methanocellales*, *Methanomicrobiales*, *Methanonatronaechaeales*, *Methanosarciniales*and *Methanomassiliicoccales* [19, 20]. Some uncultured lineages, e.g., *Ca.* Bathyarchaeota, *Ca.* Methanofastidiosa and *Ca.* Verstraetearchaeota, are speculated to also carry out methanogenesis based on the coding potential of their metagenome-assembled genomes [20]. The evidence was recently found in *Methanosuratincola* (formerly affiliated with *Ca.* Verstraetearchaeota) within Thermoproteota [21, 22], and even in Korarchaeia [23]. Methanogens consist of three main metabolic groups: acetoclastic, hydrogenotrophic and methylotrophic methanogens, utilizing acetate, H_2_/CO_2_ and formate, and methanol/methylamines/methyl-sulfides/ethanol as their substrates, respectively [24]. These pathways are the major players in methanogenesis in soils, despite recent findings on two novel methanogenesis pathways namely methoxydotrophy [25] and alkylotrophy [26].

In permafrost environments, 20%-60% of the microbial-source CH_4_ can be consumed by methanotrophs before emitting to the atmosphere [27, 28]. Methanotrophs consist of aerobic methane oxidizing bacteria (MOB) within Alphaproteobacteria, Gammaproteobacteria and Verrucomicrobia, as well as anaerobic bacteria *Methylomirabilales* and anaerobic archaeal methanotroph (ANME) groups within *Methanosarcinales* [24, 29]. Aerobic MOB of Alpha- and Gamma-proteobacteria are classified as type-II and type-I methanotrophs, respectively, and they are the major methane oxidizers occurring in most environments [29]. While these MOB mostly oxidize the microbial-source CH_4_ in soils and sediments [30], some groups are able to utilize atmospheric CH_4_ [31]. These atmospheric MOB have a high-affinity particulate methane monooxygenase and can utilize CH_4_ in a low-CH_4_-concentration environment, e.g., the atmosphere [32]. They have been detected in many different soils [33–36]; however, so far, only one strain, *Methylocapsa gorgona* MG08, has been isolated which was also detected in Arctic regions [31]. The atmospheric MOB are gaining increasing attentions as they are responsible for the mitigation of atmospheric CH_4_, which can help to counteract climate change.

The diversity and abundance of methane-cycling microbiomes are expected to vary across the Arctic due to the heterogeneity in the structure and physicochemical conditions of permafrost soil regions [37]. In soils, both methanogens and methanotrophs are important in determining CH_4_ fluxes [38–40]. It is therefore pivotal to characterize these microbiomes across space and time, for a fundamental understanding of players in determining CH_4_ dynamics in permafrost soils. However, due to the reduced accessibility of the Arctic permafrost region for sampling, the diversity of these microbiomes and the dominant microbes in these soils are poorly characterized, particularly across horizons and on a broader, pan-Arctic scale. Moreover, it remains unclear how different water conditions resulting from permafrost thaw impact the methane-cycling microbiome abundance and composition.

This study aims to reveal the pan-Arctic distributions of methanogens and methanotrophs in intact permafrost soils and their future development in response to different water conditions after permafrost thaw. We analyzed the microbiomes in 621 intact permafrost soil samples, taken from eight locations across the pan-Arctic over a course of eight years (Fig. 1a) from different horizons (Fig. 1b). The relative abundance and distribution patterns of phylotypes associated with methanogens and methanotrophs across space and horizons are shown. To understand the impact of permafrost degradation on those methane-cycling microbiomes in times of climate change, we selected three hydrologically different degraded permafrost sites and their corresponding intact site in Fairbanks, Alaska. The relative abundance and community composition of methanogens and methanotrophs were compared among the degraded sites and between the degraded sites and the intact site.

**Fig. 1.**
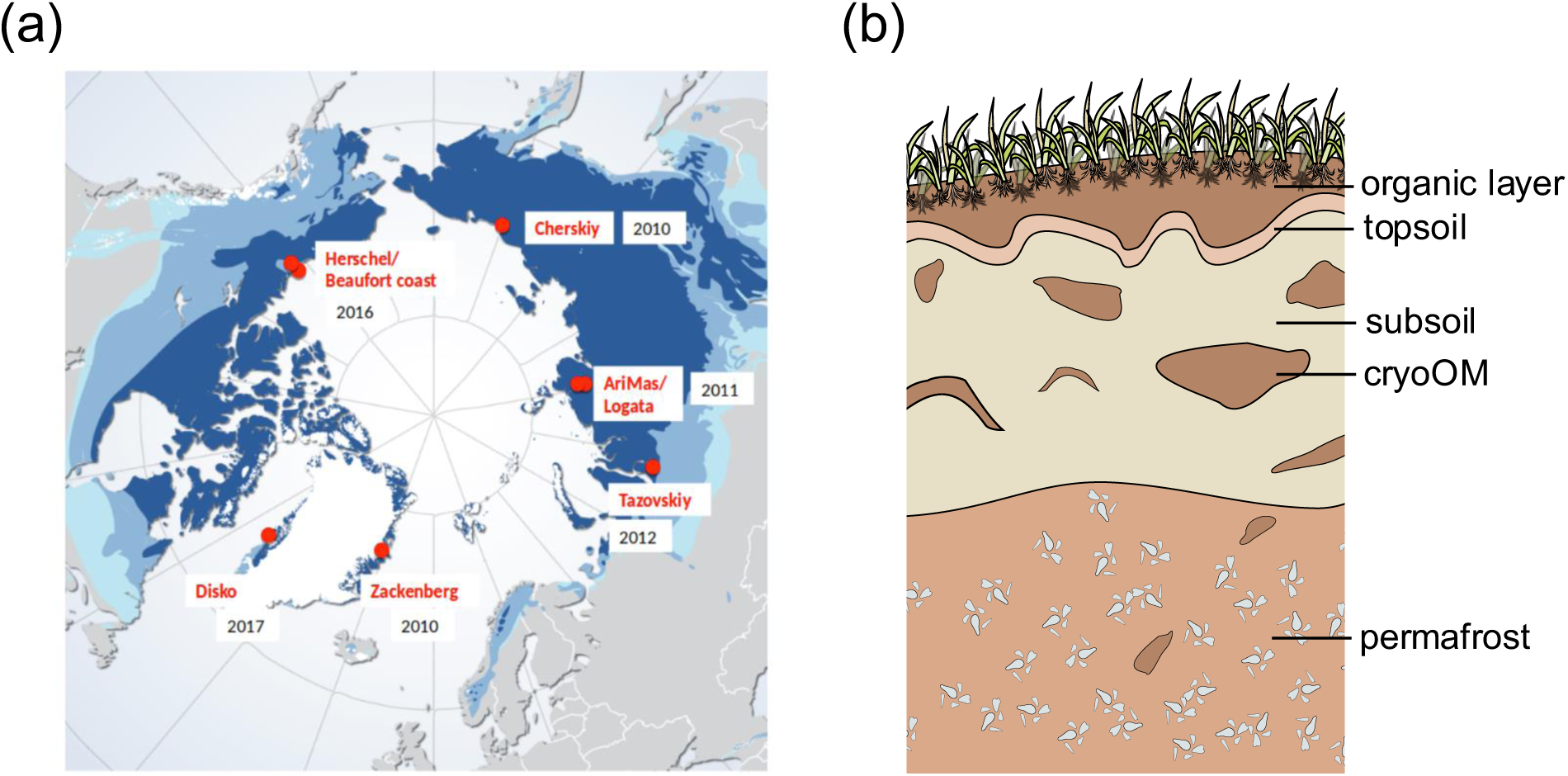
Sampling locations including (from west to east) Herschel Island and Beaufort coast (Canada), Disko Island and Zackenberg (Greenland), as well as Tazovskiy Peninsula, AriMas, Logata and Cherskiy (Russia) (a), schematic diagram of soil horizons including organic layer, topsoil, subsoil, subsoil cryoturbated organic matter (cryoOM) and permafrost (b). The base map in (a) is from https://www.grida.no/resources/5234. Dark blue refers to continuous permafrost >90% area coverage; medium blue refers to discontinuous/sporadic 10-90% coverage; light blue refers to isolated patches.

## Materials and methods

### Studying sites and soil sampling

Over the course of 8 years, soils samples were taken from 8 different intact permafrost sites around the north pole (Fig. 1a), including Herschel Island (2016) and Beaufort Coast (2016) in Canada, Disko Island (2017) and Zackenberg (2010) in Greenland, and Logata (2011), AriMas (2011), Tazovskiy (2012) and Cherskiy (2010) in Siberia, Russia. For each sampling, soil samples were taken from different horizons, including organic layer, topsoil, subsoil, cryoturbated organic matter (cryoOM) and permafrost (Fig. 1b). The sampling campaigns were conducted within different projects. The detailed descriptions of these sites and sampling protocols are available in [41–44]. The number of samples taken at each layer in each site is shown in Fig. S2 and S3. In total, 621 samples were collected.

A ninth intact permafrost site is located in the city of Fairbanks, in the Interior Alaska, USA (Fig. 2a). This Intact permafrost site (64° 51′ 56.1′′ N, 147° 51′ 18.9′′ W) is close to Smith Lake with a shallow permafrost table and no major degradation (Fig. 2b), similar to other intact sites from the pan-Arctic (Fig. 1a). To study the impact of different water conditions on microbiomes after permafrost thaw, two degraded permafrost sites in the vicinity of the Intact site were selected. The Wet degraded permafrost site (64° 52′ 02.4′′ N, 147° 51′ 17.3′′ W) is located on a low hill at the bottom of the depression with degraded permafrost soils and high-water contents (Fig. 2b). The Dry degraded permafrost site (64° 51′ 33.5′′ N, 147° 51′ 18.0′′ W) is at the shoulder of a north-facing slope with well-aerated and degraded permafrost soils and low water contents (Fig. 2b). A detailed description of these three sites is available in [45]. Additionally, a third degraded permafrost site was investigated on a mid-slope position (64° 51’ 41” N, 147° 51’ 33” W) with soil development and moisture conditions being in between those of Dry and Wet sites (Fig. 2b), in the following referred to as Intermediate-Wet site.

**Fig. 2.**
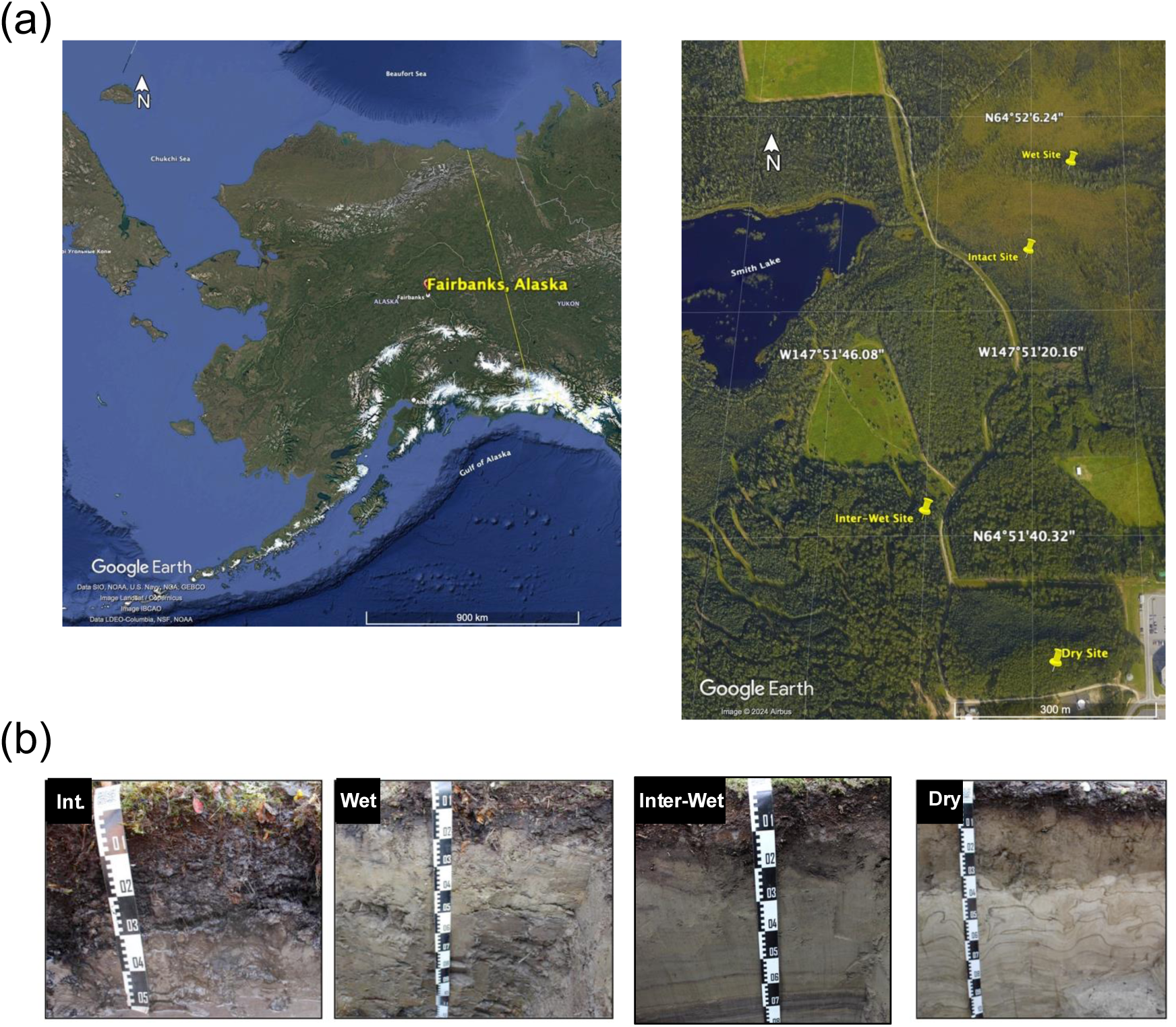
Sampling sites and their soil profiles associated in Fairbanks, Alaska (a), and the soil profiles of the four sites with different water conditions (b). Int., Intact; Inter-Wet, Intermediate-Wet. This figure is modified from [45].

Soil samples in Alaska were taken in late August-early September in 2019 and 2021. In 2019, each three soil pits (150 × 100 cm) were dug with a depth of either ∼100 cm for degraded (Dry and Intermediate-Wet) sites or until reaching the permafrost surface for the non-degraded (Intact) site. In 2021, we replaced the Intermediate-Wet degraded site with a Wet degraded site as described above, for a better comparison between distinct hydrologically degraded sites. Only one pit was dug at each site and the same approach was used for sampling the profile as in 2019. In compensation, we additionally sampled soils from 6 satellite pits at the depths of top- and sub-soil layers (∼5 cm or ∼50 cm, respectively) to account for the large, small scale heterogeneity of the soils in this area. The number of samples taken from each different depth is shown in Fig. S4.

### Soil physicochemical properties

The soil water content and pH were measured for samples taken from the 8 pan-Arctic sites (Fig. 1a). Water content was measured as water to fresh soil weight ratio by drying the soil at 105 °C until weight stabilized. The pH values were measured using a soil-water-suspension with a fixed 5:2 ratio (water to soil). The water content of Disko samples and the pH of Zackenberg samples were not available due to different managements and measurements of the involving projects.

Soil samples taken in Alaska were analyzed on a variety of soil physicochemical parameters, including soil moisture, pH, soil organic carbon (SOC), total carbon (TC) and nitrogen (TN), base saturation (BS), dissolved organic carbon (DOC) and nitrogen (DN), micro- and macro-aggregate (MiA and MaA) proportions, and organic carbon and nitrogen in MiA and MaA. Data were adapted from [45].

### Microbiome analysis

Soil samples taken from the 8 pan-Arctic sites (Fig. 1a) were processed with DNA extraction and 16S rRNA gene amplicon sequencing using the primer pair 515F/806R. Illumina sequencing with pair-end 100 bp was performed with Cherskiy and Zackenberg samples collected in 2010, while the other samples were sequenced with a later pair-end 250 bp Illumina platform. To make all sites comparable, sequences were all trimmed to 100 bp for downstream analysis. The UNOISE algorithm [46] was used to denoise and error-correct the sequencing data, and zero-radius Operational Taxonomic Units (ZOTUs) were generated. The taxonomy of each ZOTU was assigned against a modified SILVA128 database [47] using lowest common ancestor algorithm with MEGAN5 [48]. Based on 621 analyzed datasets, a subset of ZOTUs associated with methanogens and methanotrophs (Table S1) were used for downstream analyses. The core members that existed in all sites except for Tazovskiy (due to absence or low abundances of both methanogens and methanotrophs) were identified. The representative sequences of associated ZOTUs were further used as enquiry for NCBI BLASTN against the 16S rRNA sequence (bacteria and archaea) database to verify the taxonomy identification and, if possible, acquire species information.

Alaska soil samples (Fig. 2a) in 2019 were transported on ice and kept at −20 °C before the DNA extraction. DNA was co-extracted with the RNA using the RNeasy PowerSoil total RNA kit and the RNeasy PowerSoil DNA elution kit (Qiagen, Hilden, Germany) from ∼2 g of homogenized soil samples. In 2021, approximately 3 g of each soil was mixed with 2 volumes of LifeGuard Soil Preservation Solution (Qiagen, Hilden, Germany) directly in the field. All treated soils were kept at 4 °C before the DNA extraction. DNA was also co-extracted with the RNA using the same kits. The LifeGuard solution was removed by centrifugation and ∼2 g of the treated soil was used for the extraction. The extracted DNA samples were used for 16S rRNA amplicon sequencing using Illumina MiSeq platform (pair-end 250 bp) using the primer pair 515F/806R. The data in Alaska samples were analyzed separately and the phylotypes of methanogens and methanotrophs were inferred as amplicon-sequencing-variants (ASVs). However, ASVs and ZOTUs both provide species-level resolved phylotypes [46, 49] and are thus comparable with each other.

The *dada2* pipeline [49] was used to process the Alaska data in R v3.6.3 [50]. Sequences failing to meet the filter criteria (maxEE = 2, truncQ = 2, maxN = 0) were removed. Those filtered sequences were de-replicated, the ASVs were deduced, and the paired-end sequences were merged. Afterwards the chimeric sequences were removed. The sequence of each ASV was assigned to taxonomy using the SILVA 138.1 database. ASVs associated with methanogens and methanotrophs were identified with the same criteria (Table S1) but considering changes in some taxa names or hierarchical placements in the new database [51]. Those ASV sequences were also used as query to search for the closely related species on NCBI as described above.

The representative sequences of methanotroph and methanogen ASVs and their closely related reference sequences were used for constructing a phylogenetic tree. To further classify *Methylobacter tundripaludum* associated ASVs, the 16S rRNA genes from metagenome-assembled genomes (MAGs) of several *M. tundripaludum* species were also included. All sequences were aligned using SINA Aligner [52] with denovo mode. The aligned sequences were trimmed to the same length using MEGAX [53]. The trimmed alignment was used as input for phylogenetic tree construction using IQ-TREE [54]. The model for a maximum likelihood tree was determined using the Tree Inference function in IQ-TREE with 1000 bootstraps. FigTree v1.4.4 was used to visualize the tree.

To link the methanotroph ASVs in Alaska to ZOTUs in the pan-Arctic, a local BLASTN was run using ASVs as the enquiry and ZOTUs as the database. Only ASVs showing a 100% identity to a ZOTU are considered as the potential same phylotype.

### Statistics

The downstream analyses were done with R v3.6.3 [50]. The normality of abundance data distributions was checked before linear regressions or t tests using the Shapiro-Wilk test. If not normally distributed, the data were log_10_ transformed. A minimum value was added to avoid zeros when necessary, before the transformation. The coefficients and significance level of the correlations were determined by Pearson’s correlation analysis using *vegan* v2.5.6 package [55].

The significant correlations were further confirmed by non-parametric Spearman’s correlation analysis using *vegan*. Pairwise t tests were performed to compare the means of methanogen and methanotroph relative abundances between each two sites or horizons. Spearman’s correlation was also used to check the correlations between methanogen and methanotroph abundances and environmental parameters for Alaska samples. For all multiple comparisons, *P* values were adjusted by the false discovery rate (FDR) method. All of the plots were generated using *ggplot2* v3.3.3 package [56].

## Results and discussion

### Functional guild abundances across the pan-Arctic and soil horizons

Methanogen relative abundance (in total microbiome) varied strongly between locations, from the highest found in Herschel Island (∼0.5%) and Zackenberg (∼0.75%) to almost undetected in Disko Island and Tazovskiy (Fig. 1a). The methanogen distribution along the vertical profiles was often location-specific, sometimes increasing with depth, and sometimes having the highest abundance in the organic layers (Fig.1c and S2). Nevertheless, methanogen relative abundance and soil water content were significantly positively correlated (Fig. S1a), indicating that the hydrologic status is a key driving factor for methanogen abundance by limiting oxygen diffusion into the soil profile. pH showed no significant correlation with methanogen relative abundances (Fig. S1b).

Methanotroph relative abundance (in total microbiome) followed the methanogen pattern, with the highest (∼0.32%) and lowest (almost undetected) abundances found in Zackenberg and Tazovskiy, respectively (Fig. 1b). Along the soil profiles, methanotrophs were less abundant in the organic layer and topsoil compared to the other deeper layers (Fig. 1d), but this pattern again varied between sites (Fig. S3). Neither water content nor pH showed a significant correlation with methanotroph relative abundance (Fig. S1c and S1d). However, methanotroph abundances were significantly positively correlated with methanogen abundances (Fig. S1e), demonstrating that methanotrophs rely on the CH_4_ produced by the methanogens, and thus that their abundance is predominantly driven by the resource availability rather than abiotic factors such as water or pH.

### Functional guild compositions across the pan-Arctic and soil horizons

Methanogens conducting hydrogenotrophic methanogenesis dominated the methane producing community (Fig. 4a). We considered phylotypes found in all locations (excluding Tazovskiy) as members of the pan-arctic core methanogenic microbiome. One member was associated with the uncultured candidatus family *Methanoflorentaceae* (former Rice-cluster-II) which had been found prevalently in thawing permafrost [57, 58]. Another phylotype was associated with the H_2_-dependent methylotrophic order *Methanomassilicoccales*, more specifically with the currently uncultured wetland cluster widely distributed in peatlands [59]. Another one was closely related with *Methanosarcina lacustris* isolated from anoxic lake sediments [60] and cold terrestrial habitats [61], that can use multiple methanogenesis pathways. The most abundant phylotype, however, was closely related to *Methanobacterium lacus* [62, 63], which can perform methanogenesis from both H_2_/CO_2_ and methanol/H_2_ pathways.

The pan-arctic methanotrophs were dominated by type I methane oxidizing bacteria (MOB, Fig. 4b). Nevertheless, one of the two phylotypes of the *Ca.* Methanoperedens clade (formerly ANME-2d), was notably abundant across many locations, especially in the subsoil (Fig. 4b), suggesting a potential vital role of anaerobic methane oxidation in mitigating methane emissions in anoxic deeper layers of permafrost soils. Among the pan-Arctic MOBs, only one was from type II, related to *Methylocella palustris*, a facultative methanotroph first isolated from peat bogs [64]. In contrast, six were closely related to the obligate type I methanotroph *Methylobacter tundripaludum* SV96 isolated from Svalbard peatlands [65]. Taken together, these six *M. tundripaludum* phylotypes accounted for in average 68% of the total methanotrophic community. This proportion varied across studied locations, ranging from min. 28% (Tazovskiy, Russia) to max. 98% (Disko, Greenland). This finding is remarkable, since it suggests that the microbial CH_4_ filter in permafrost-affected soils across the Arctic is of a strikingly low diversity and dominated by one single species. Supporting this, *M. tundripaludum* was also found as dominant methanotrophs in other Arctic locations, e.g., Svalbard [66] and the Lena Delta in Siberia [67]. A recent study showed physiological adjustments of *M. tundripaludum* to varying temperatures [68], which might explain the prevalence and dominance of *M. tundripaludum* across heterogeneous Arctic regions.

### Responses to different water status after permafrost degradation – case study in Alaska

The relative abundance of methanogens in the Intact site was < 0.01% in both 2019 and 2021 (Fig. S4a), which is among the lowest relative abundances detected in the other intact sites across the pan-Arctic (Fig. 3a). Accordingly, only 1 phylotype closely related with *Methanobacterium lacus* was found in the Intact site in 2021 (Fig. 5a). However, 4 phylotypes associated with *Methanomassiliicoccus luminyensis* (Fig. S5) were mainly found under Dry and Intermediate-Wet conditions (Fig. 5a), but still only accounting for up to 0.03% of the total microbiome (Fig. S4a). They were only detected in the deeper soil layers (>30 cm below the surface) (Fig. 5a).

**Fig. 3.**
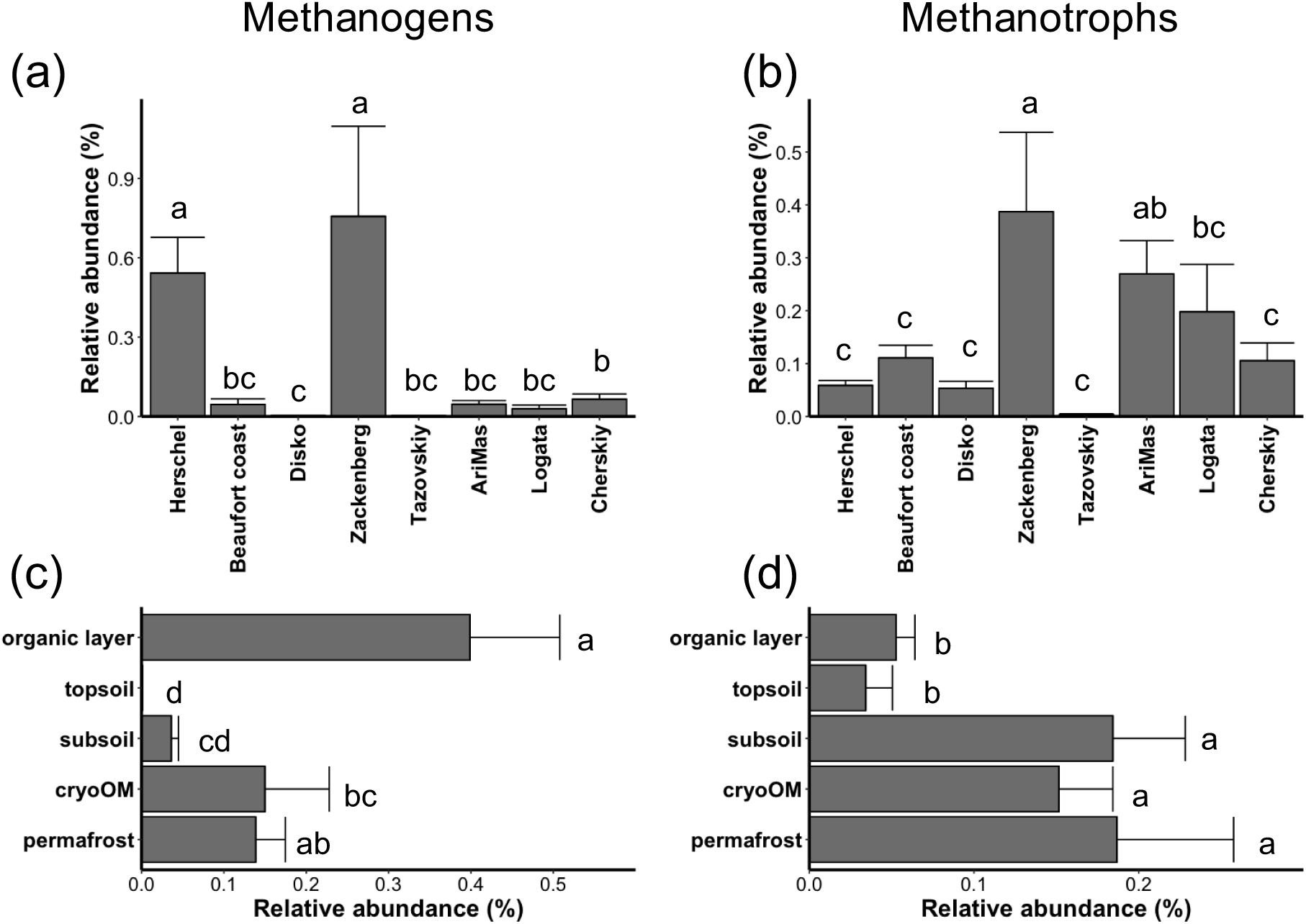
Relative abundances of methanogens (a) and methanotrophs (b) in different locations, and relative abundances of methanogens (c) and methanotrophs (d) in different horizons. Abundances are shown as mean + standard error. The bars with no common letter are significantly different (*P* < 0.05) characterized by Kruskal-Wallis *post hoc* Dunn’s tests with *P* values adjusted by the false discovery rate method.

The methanotrophs were less diverse in the Intact site compared to the other pan-Arctic sites (Fig. 4b and 5b). Only *Methylobacter* phylotypes were detected in 2019, with additionally two *Crenothrix* ASVs detected in 2021. Their relative abundance in the Intact site is comparable to that in Herschel Island and Disko Island (Fig. 3b and S4b). In contrast, the methanotrophic communities in Dry and Wet sites were dominated only by *Methylocapasa* and *Methylobacter*, respectively (Fig. 5b). In the Intermediate-Wet site, both groups were found, with additionally *Crenothrix* and *Ca*. Methylomirabilis phylotypes detected (Fig. 5b). Compared to the Intact site, the relative abundance of methanotroph was higher in the Dry and Intermediate sites and similar in the Wet site (Fig. S4b). All type I methanotrophs (MOB) were only found in depth >30 cm below the surface where methanogens were detected, while the type II methanotrophs (MOB) in the Dry site were mostly found in upper soil layers, with a few found in the deeper layers (Fig. 5a and 5b).

**Fig. 4.**
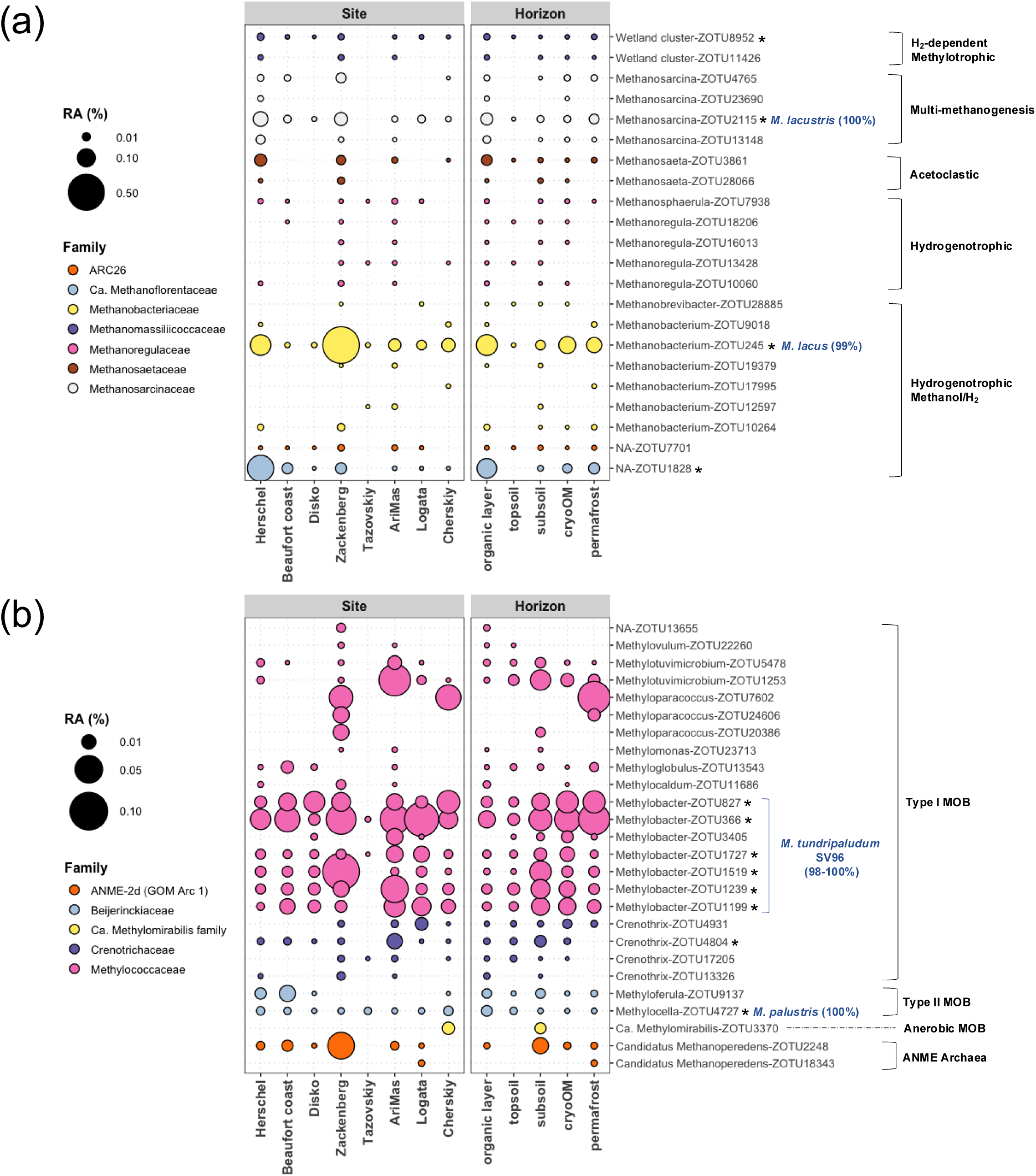
The relative abundance of each ZOTU associated with methanogens (a) or methanotrophs (b) in different sites and horizons. RA, relative abundance; NA, not assigned at genus level; MOB, methane oxidizing bacteria; ANME, anerobic methanotroph; *, core methanogen or methanotroph members that were found in all sites (excluding Tazovskiy). Core members showing a match with >97% identity to sequences in NCBI 16S rRNA gene sequence (bacteria and archaea) database are additionally labelled with the species name and their identity value to that species.

The distributing pattern of methanotrophs in different Alaska sites is likely due to the different water conditions. In the Dry site, the *Methylocapasa* ASVs are closely associated with *M. palsarum* and *M. gorgona* (>97% identity). It is known that *Methylocapasa* hosts atmospheric MOB and that *M. gorgona* MG08 is so far the only isolated atmospheric MOB [31]. Due to their high-affinity to CH_4_, these MOB can utilize CH_4_ in the atmosphere [32]. This explains why the Dry site hosted only *Methylocapasa* MOB. Dry conditions create aerobic environments which attenuate anaerobic methanogenesis, and the lack of microbial CH_4_ source further hampers the “conventional” but not the atmospheric MOB. In contrast, the Intact and Wet sites only hosted “conventional” type I MOB in deeper layers due to wetter and more anaerobic conditions under which methanogens could survive and potentially produce CH_4_. In the Intermediate-Wet site, the dominant MOB switched from *Methylocapasa* in top layers to *Ca*. Methylomirabilis in middle layers then to *Methylobacter* and *Crenothrix* in deep layers (Fig. 5b). The anaerobic and type I MOB found in the middle and deeper layers were also likely supported by the methanogens living in these layers (Fig. 5a). The existence of both atmospheric and “conventional” MOB in turn indicated an intermediate water condition providing niches for both groups in the Intermediate-Wet site.

**Fig. 5.**
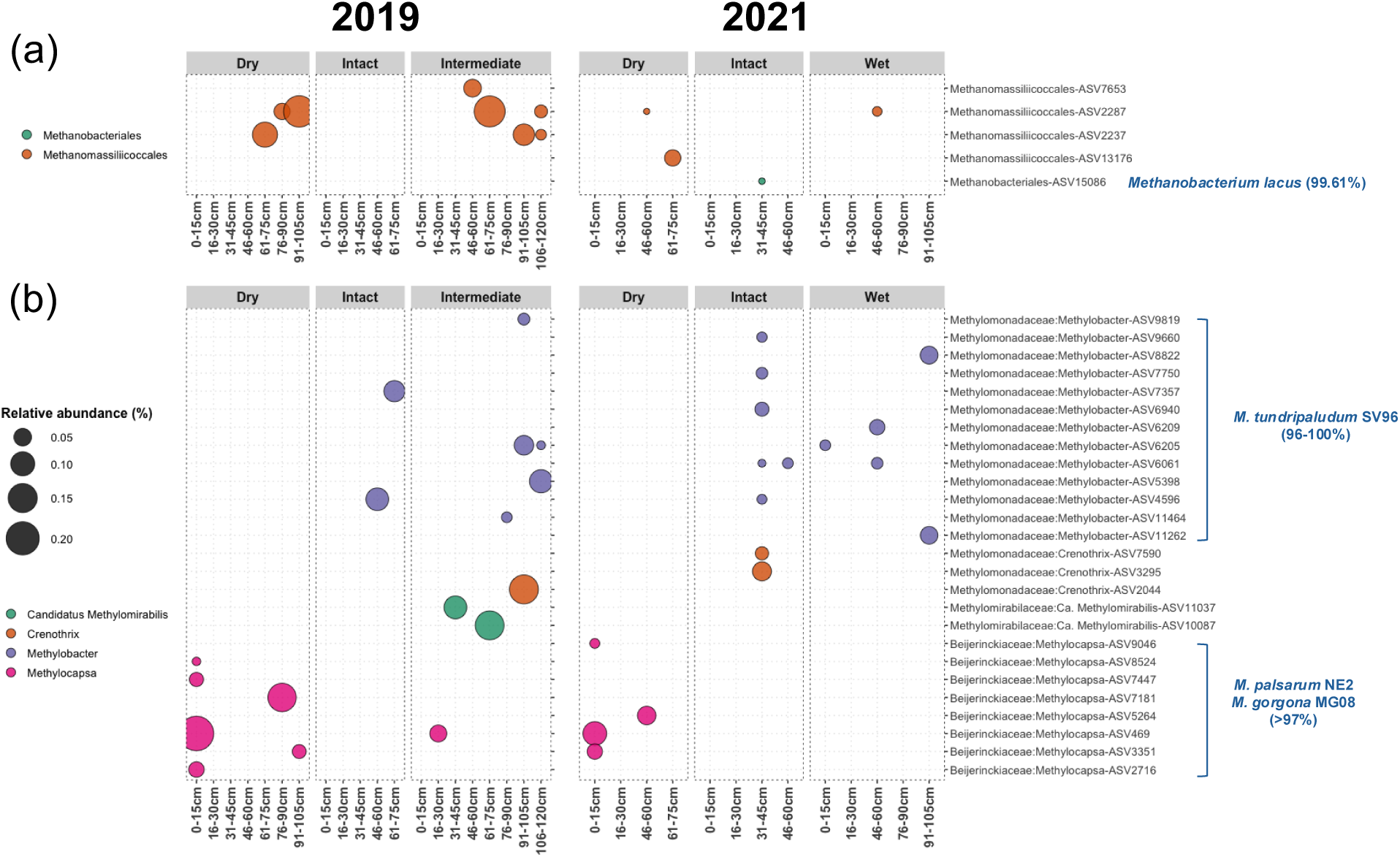
The relative abundance of each ASV associated with methanogens (a) or methanotrophs (b) in different scenarios and depths in 2019 and 2021 in Fairbanks, Alaska. The best matches (>97% identity) of ASVs to sequences in NCBI 16S rRNA gene sequence (bacteria and archaea) database are shown as the matched species name and their identity value to that species.

These distribution patterns suggested that type I MOB found in these sites were dependent on the methanogens as their main CH_4_ suppliers. This is supported by their congruent distributions along the depth (Fig. 5) as well as their congruent responses to the soil physiochemical properties (Fig. 6). Both methanogens and type I MOB showed a positive correlation with depth, pH and MiA%, and a negative correlation with C/N and temperature, although the correlations were mostly only significant for samples collected in 2019 (Fig. 6a and 6b). In contrary, type II MOB positively correlated with SOC, total TN, and the C/N ratio, but negatively with pH. This suggested that their proportion in the community increased with higher soil C and significantly increased in acidic conditions. Due to their ability to acquire CH_4_ from the atmosphere, we argue that they can contribute to C input to the drained permafrost soil. This is evidenced by their positive correlation with SOC, as they convert CH_4_ to CO_2_, which can then be directly used by autotrophs to fix it and become part of the SOC. Recent study of Zheng et al., 2024 [69] found an identical relationship of type II MOB with SOC in paddy soils. This is a crucial finding because the atmospheric CH_4_ fixation by increased number of type II MOB to SOC via autotrophic metabolism may contribute in the future to mitigation of atmospheric CH_4_ in drier landscapes after permafrost degradation.

**Fig. 6.**
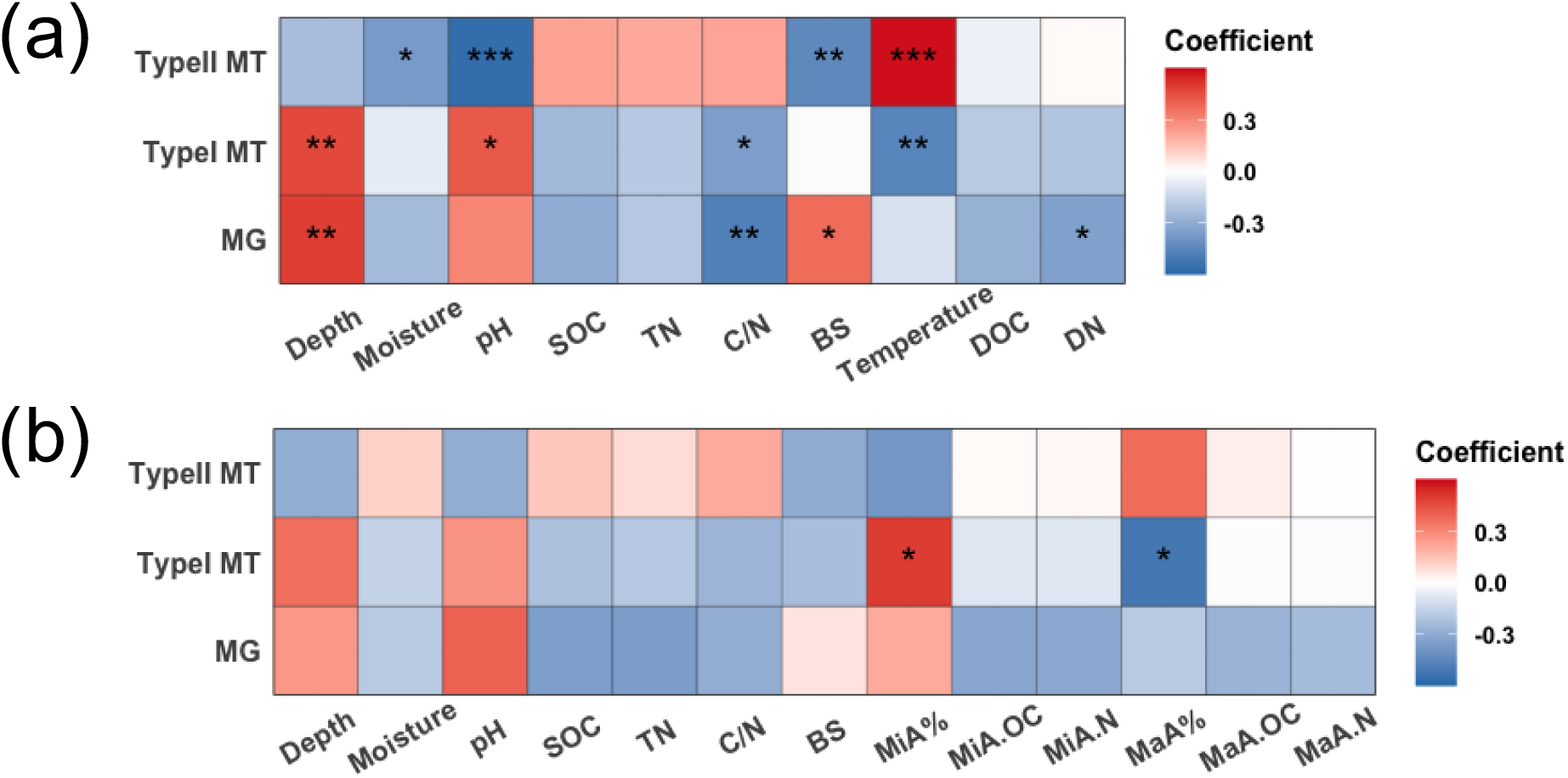
Spearman’s correlations between relative abundance of methanogens (MG) and methanotrophs (MT) and soil physiochemical properties in 2019 (a) and 2021 (b) in Alaska samples. Asterisks indicate adjusted *P* values, \**P* < 0.05, \*\**P* < 0.01, \*\*\**P* < 0.001. SOC, soil organic carbon. TN, total nitrogen. BS, base saturation. MiA, micro-aggregation. MaA, macro-aggregation.

### Identification of *M. tundripaludum* phylotypes found in Alaska

Similar to the pan-Arctic sites, *M. tundripaludum* SV96 associated phylotypes dominated the MOB community in the Intact and Wet sites (Fig. 5b). This is astonishing as their dominance is consistent across the pan-Arctic permafrost affected soils, even including the Wet degraded permafrost site. It is noteworthy that SV96 is the only available *M. tundripaludum* strain found in the NCBI 16S rRNA gene sequence (bacteria and archaea) database. However, *M. tundripaludum* is a diverse group hosting different strains based on MAGs [70]. To identify whether these *M. tundripaludum* associated ASVs were exclusively related with SV96, we constructed a phylogenetic tree by including several 16S rRNA gene sequences from *M. tundripaludum* MAGs for further classification (Fig. 7). Interestingly, all the *Methylobacter* and *Crenothrix* phylotypes, all different *M. tundripaludum* strains, as well as *M. psychrophilus* Z-0021 and *M. psychrotolerans* Sph1, formed a single cluster (Fig. 7). It is known that *Crenothrix*, *M. psychrophilus* Z-0021 and *M. psychrotolerans* Sph1 are the most closely related species of *M. tundripaludum* based on both 16S rRNA gene and *pmoA* (marker gene of methanotrophs) [70]. The detected sequence length in our study is 250 bp and could not hint a full-length 16S rRNA gene-based phylogeny. Therefore, the lack of information from the missing parts of 16S rRNA gene might lead to the misplacement of closely related species, i.e., *M. psychrophilus* Z-0021 and *M. psychrotolerans* Sph1. The *Crenothrix* ASVs were even more closely clustered with some *M. tundripaludum* than some *Methylobacter* ASVs (Fig. 7), suggesting that these ASVs might also be *M. tundripaludum* related.

**Fig. 7.**
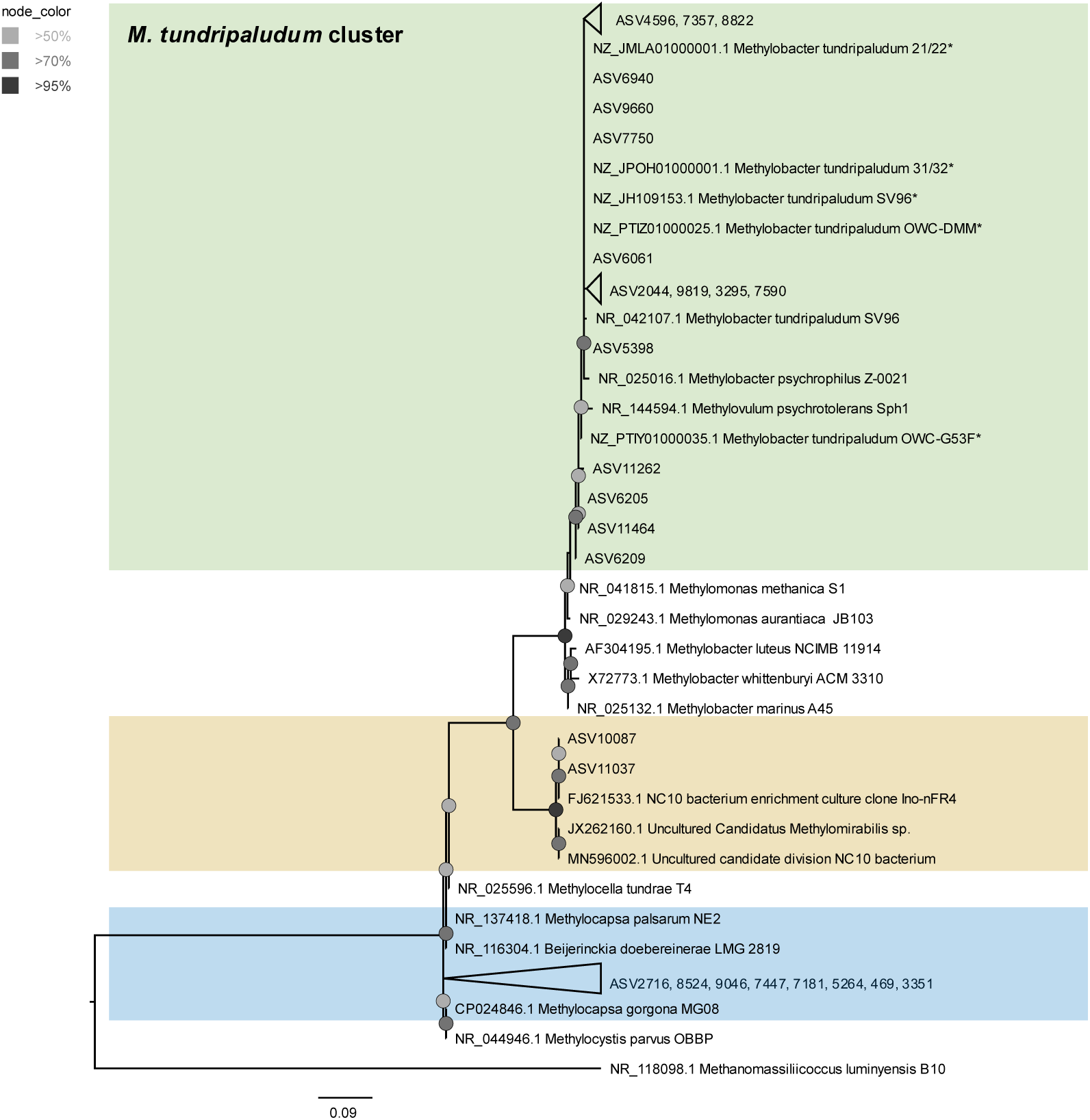
Phylogenetic tree inferred from 16S rRNA gene sequences of methanotrophic ASVs in Alaska samples and their closely related sequences found in NCBI 16S rRNA gene sequence (bacteria and archaea) database. *Several 16S rRNA genes from metagenome-assembled genomes of *M. tundripaludum* are included to distinguish different *M. tundripaludum* associated phylotypes. The tree was inferred by maximum likelihood with IQ-TREE based on TIM3+G4+F model. The node colors indicate ultrafast bootstrap support > 50%. Background colors indicate different ASV clusters.

We ran a local BLASTN to link Alaska ASVs to the pan-Arctic ZOTUs. Most of the type I MOB ASVs could be linked to a ZOTU with 100% identity (Table S2), suggesting that most of these detected Alaska *Methylobacter* were likely identical to those found in the pan-Arctic and that these pan-Arctic *M. tundripaludum* might not be only associated with SV96 but also with other *M. tundripaludum* strains. Nonetheless, this does not alleviate the striking fact that *M. tundripaludum* as a single species dominated the microbial CH_4_ filter in permafrost-affected soils across the Arctic.

### The methane-cycling microbiome in Arctic permafrost: unlocking future climate scenarios

Our data suggested that both dry and wet conditions after permafrost thaw had an impact on the methanogen and methanotroph communities. There was a slight increase of methanogen relative abundance and the number of their phylotypes in both Dry and Wet or Intermediate-Wet sites (Fig. 5a). Despite their low proportions in microbiome in all sites (Fig. S4a), these patterns hinted that permafrost degradation might lead to an increase of methanogen richness. On the other hand, the methanotrophs showed a much bigger diversity and a completely contrasting pattern of their community compositions between the Wet and Dry sites. The single dominant group switched completely from *Methylobacter* in the Wet site to *Methylocapsa* in the Dry site (Fig. 5b). The latter is known to be closely related with atmospheric MOB, suggesting that drier conditions after permafrost degradation might even result in a CH_4_ sink as these MOB could consume CH_4_ from the atmosphere. While the status of being wet or dry after permafrost thaw depends on native factors such as vegetation, topography, ice storage and substrate [71], many thawed areas will likely experience drier conditions as a result of higher temperatures and increased evaporation in a warming climate [72]. Due to global warming, extreme weather events, such as summer drought, are happening more frequently than before [73], which may even boost these processes. Our data suggest that atmospheric MOB might play a more important role in mitigating CH_4_ emissions from drained landscapes after permafrost thaw in a future warmer climate.

## Conclusion

This is the first study to show the distribution of methanogens and methanotrophs across regions and horizons in permafrost-affected soils on a pan-Arctic scale. Methanogens were more abundant in organic and deep layers with high water contents. Methanotrophs were also more abundant in the rather oxygen-depleted deeper layers, pointing to their dependence on methanogens for the CH_4_ source. However, these patterns and their relative abundances also varied across locations. Four methanogens with a diverse set of methanogenesis pathways indicate a certain flexibility for different methanogenesis substrates and methanogenic conditions in permafrost. Most strikingly, methanotroph phylotypes closely related with *M. tundripaludum* dominated this functional guild in the pan-Arctic microbiome as well as in the wet degraded permafrost in Alaska. This indicates that *M. tundripaludum* as a single species might act as the major microbial CH_4_ filter in Arctic soils. However, under drier conditions, this group could be replaced by atmospheric MOB, which can result in more CH_4_ uptake in Arctic regions. These findings are crucial for estimating the future methane filter functioning and magnitude in Arctic tundra and taiga soils after permafrost degradation.

## Data Availability

Sequencing data from Cherskiy are available from MG-RAST (project numbers 44668685.3– 44668734.3). Sequencing data from Zackenberg, Logata and AriMas are available from the Qiime database (http://www.microbio.me/emp/, study no. 1034). Sequencing data from Disko Island, Herschel Island, Beaufort Coast and Alaska are available from European Nucleotide Archive of European Molecular Biology Laboratory (accession number PRJEB72670 for Disko Island; PRJEB38326 for Herschel Island and Beaufort Coast; PRJEB83087 for Alaska). All primary and secondary data and R codes are available as supplementary materials.

## Acknowledgements

This study was supported by Deutsche Research Foundation (DFG) project GU 406/35-1 and UR 198/4-1, and the Czech Science Foundation (GACR) project 20-21259J. The authors thank Marc Piecha for his help with the conceptual figure.

## Author contributions

**Haitao Wang**: Conceptualization; Investigation; Formal Analysis; Visualization; Writing - Original Draft Preparation. **Erik Lindemann**: Data Curation; Investigation; Formal Analysis; Visualization. **Patrick Liebmann**: Investigation; Writing - Review & Editing. **Milan Varsadiya**: Investigation; Writing - Review & Editing. **Mette Marianne Svenning**: Writing - Review & Editing. **Muhammad Waqas**: Writing - Review & Editing. **Sebastian Petters**: Investigation; Writing - Review & Editing. **Andreas Richter**: Methodology; Writing - Review & Editing. **Georg Guggenberger**: Resources; Methodology; Writing - Review & Editing. **Jiri Barta**: Methodology; Resources; Formal Analysis; Investigation; Writing - Review & Editing. **Tim Urich**: Validation; Project administration; Funding Acquisition; Supervision; Writing - Review & Editing.

