## Supporting material for "The methane-cycling microbiome in intact and degraded permafrost soils of the pan-Arctic"

**Running head**: Pan-Arctic methane-cycling microbiome

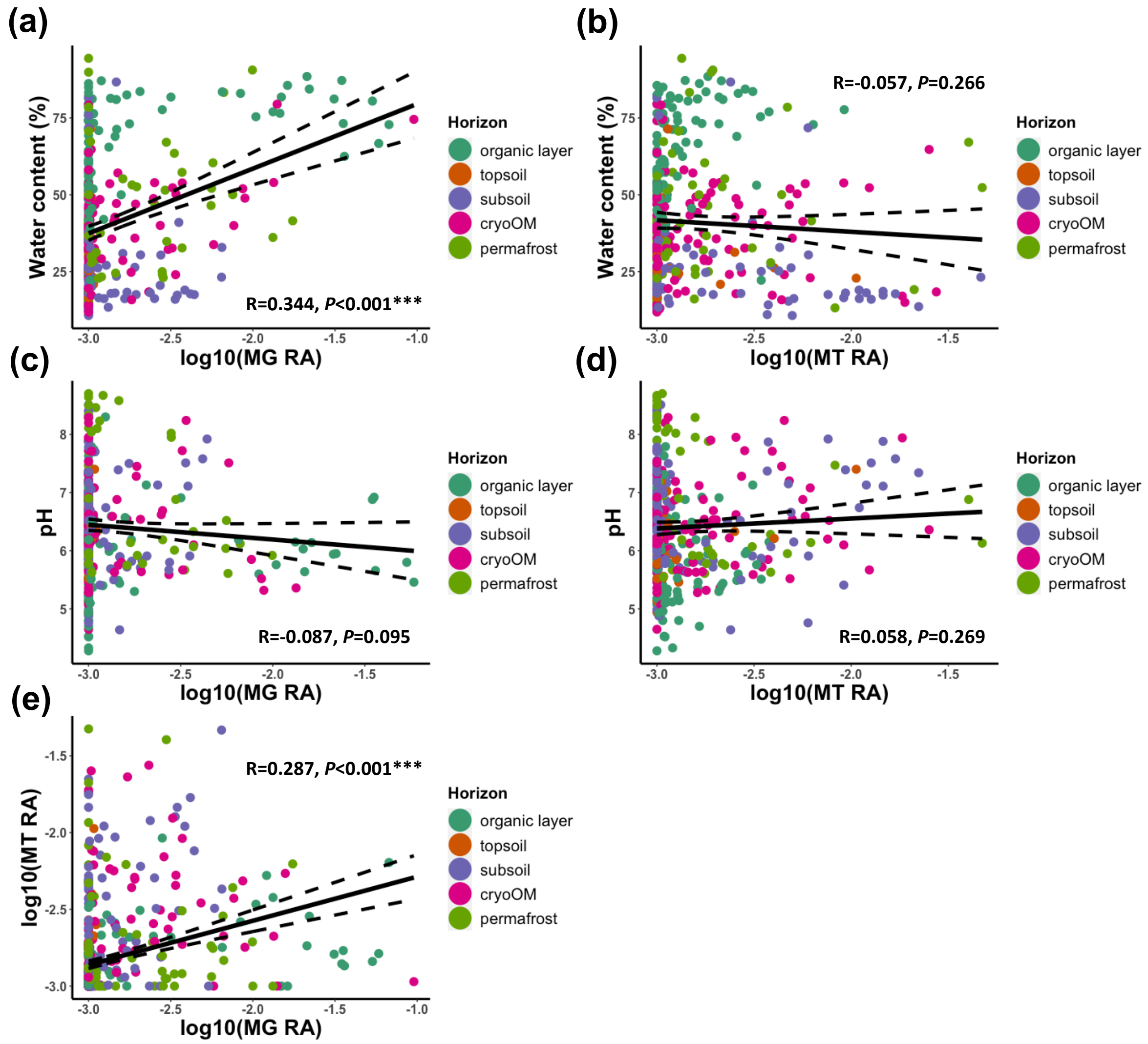


**Fig. S1** Correlations between water content and methanogen (a) and methanotroph (b) abundances, correlations between pH and methanogen (c) and methanotroph (d) abundances, and correlation between methanogen and methanotroph abundances (e). MG, methanogen; MT, methanotroph; RA, relative abundance. The R and *P* values are based on Pearson's correlation. A value of 0.001 was added to MG and MT relative abundances before $\log_{10}$ transformation to avoid zeros. The significant correlations were also confirmed as significant with Spearman's correlation.

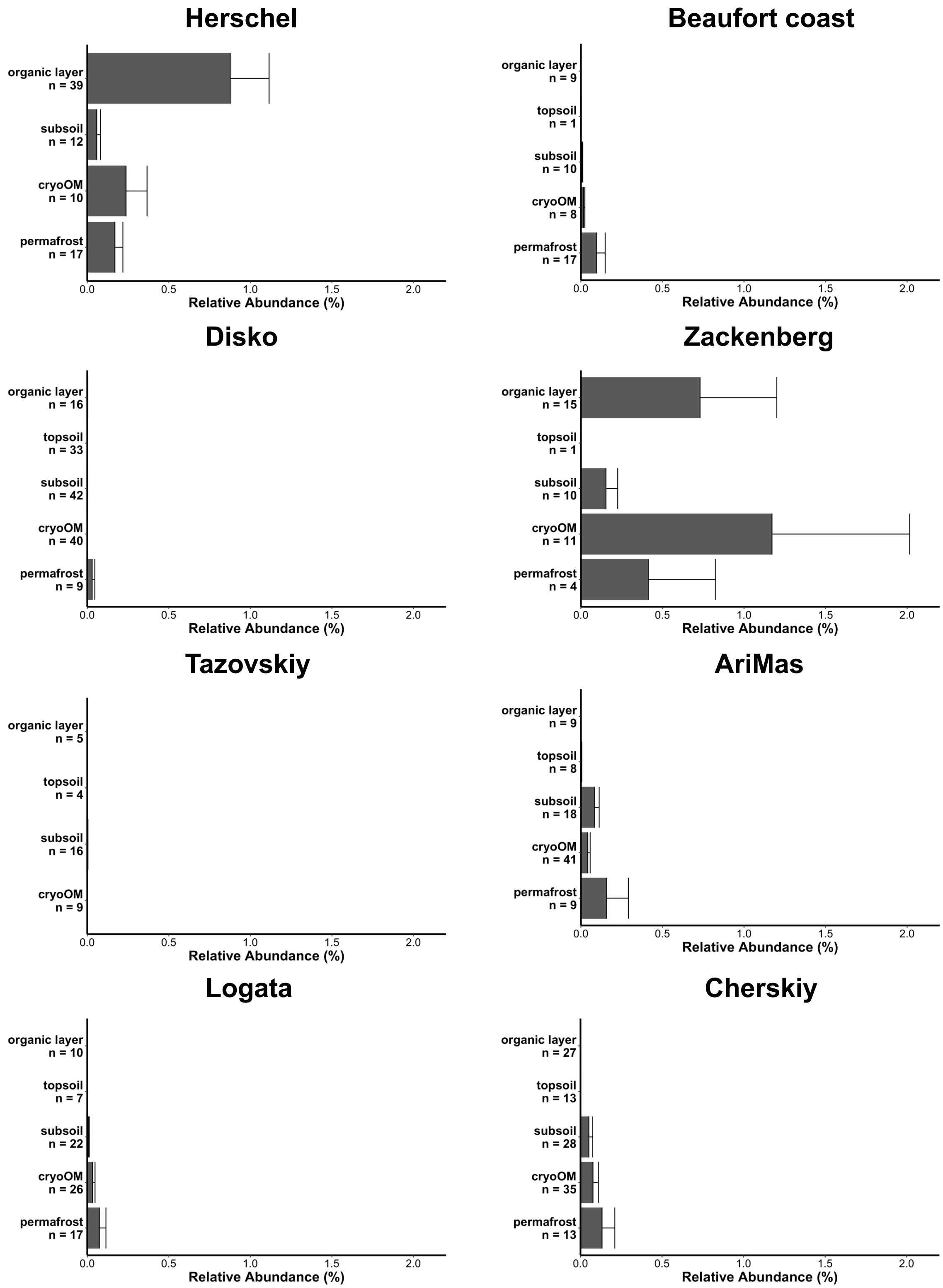


**Fig. S2** Relative abundance of methanogens in different horizons at each site. Abundances are shown as mean + standard error.

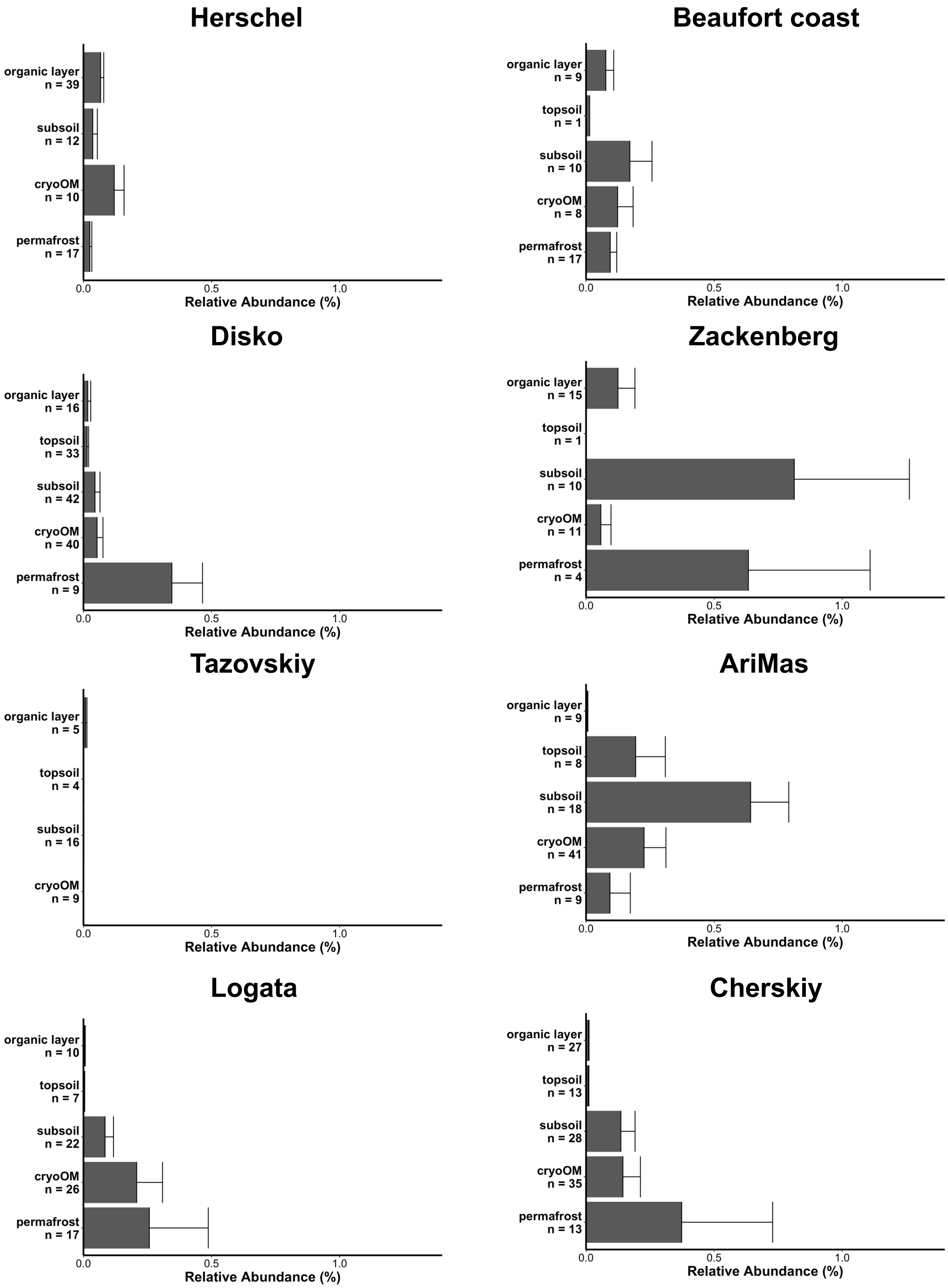


**Fig. S3** Relative abundance of methanotrophs in different horizons at each site. Abundances are shown as mean + standard error.

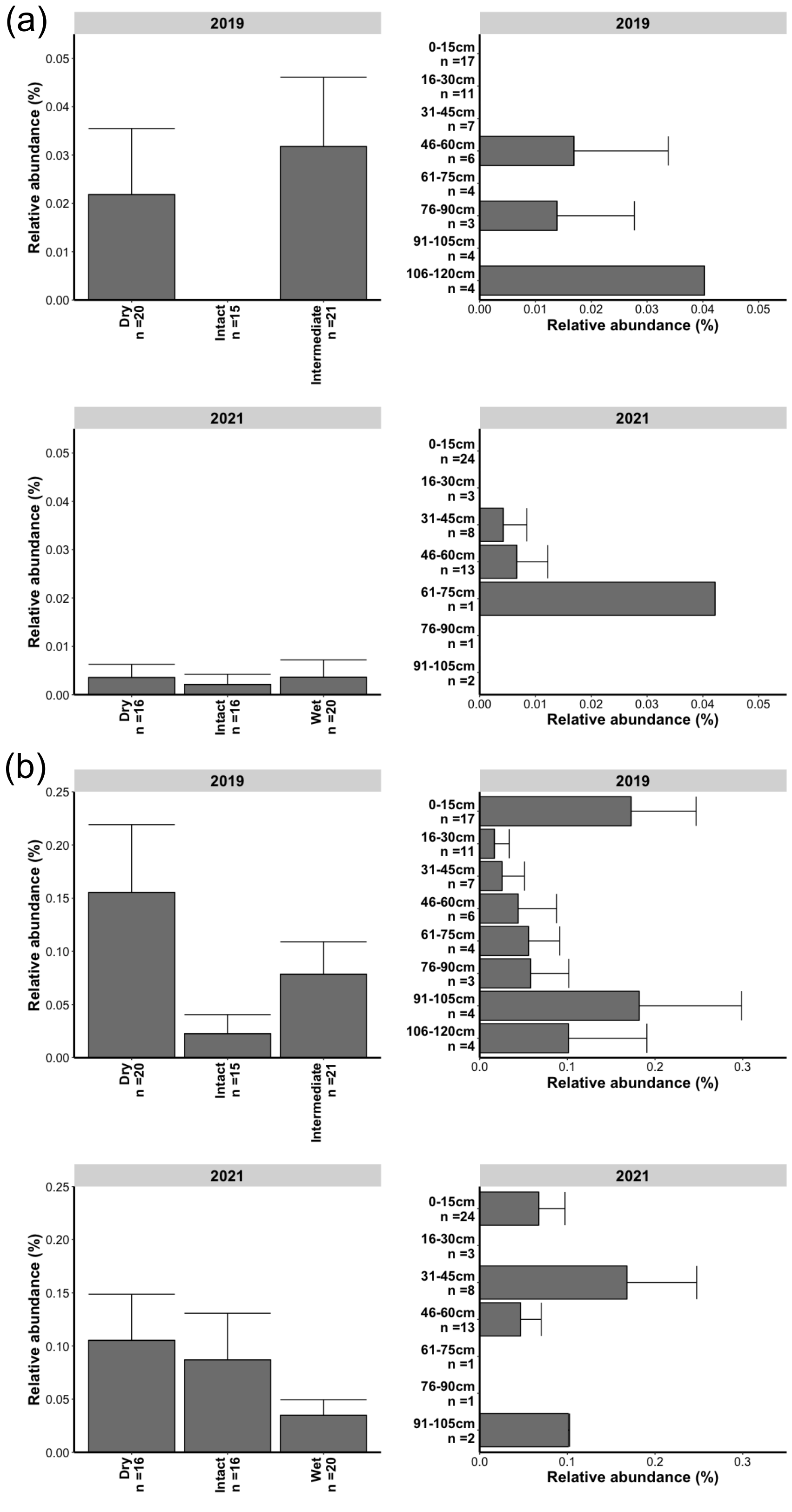


**Fig. S4** Relative abundances of methanogens (a) and methanotrophs (b) in between different scenarios and depths in 2019 and 2021. Abundances are shown as mean + standard error. The bars with no common letter are significantly different ($P < 0.05$) characterized by Kruskal-Wallis *post hoc* Dunn's tests with $P$ values adjusted by the false discovery rate method.

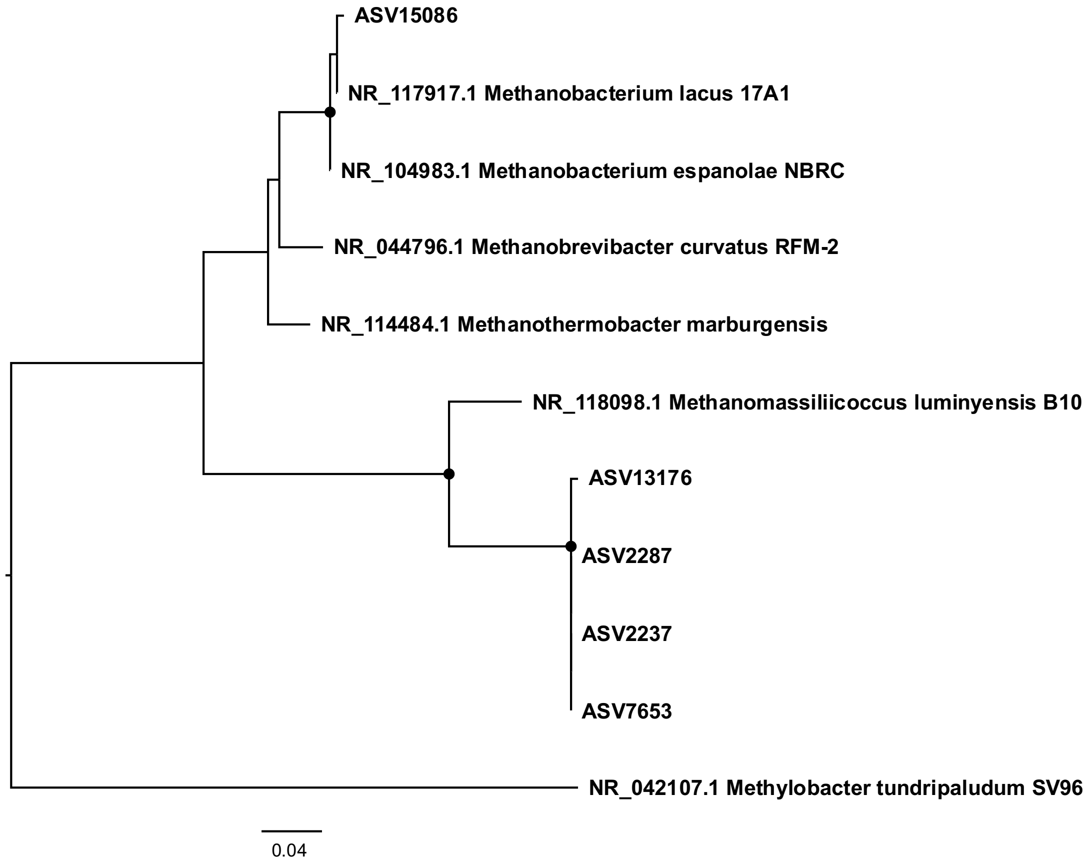


**Fig. S5** Phylogenetic tree inferred from 16S rRNA gene sequences of methanogenic ASVs in Alaska samples and their closely related sequences found in NCBI 16S rRNA gene sequence (bacteria and archaea) database. The tree was inferred by maximum likelihood with IQ-TREE based on HKY+F+I model. The node colors indicate ultrafast bootstrap support > 50%.

**Table S1** Taxonomy list used for identifying methanogens and methanotrophs

| | Phylum | Class | Order | Family | Genus | No. of ZOTUs |
|---|---|---|---|---|---|---|
| **Methanogens** | Euryarchaeota | Methanomicrobia | **Methanosarcinales** (excluding ANME) | - | - | 6 |
| | Euryarchaeota | Methanomicrobia | **Methanomicrobiales** | - | - | 7 |
| | Euryarchaeota | Methanomicrobia | **Methanocellales** | - | - | - |
| | Euryarchaeota | Methanobacteria | **Methanobacteriales** | - | - | 7 |
| | Euryarchaeota | Methanopyri | **Methanopyrales** | - | - | - |
| | Euryarchaeota | Methanococci | **Methanococcales** | | | - |
| | Euryarchaeota | Thermoplasmata | **Methanomassiliicoccales** | - | - | 2 |
| | Euryarchaeota | **Methanonatronarchaeia** | - | - | - | - |
| **Methanotrophs** | Proteobacteria | Alphaproteobacteria | Rhizobiales | Methylocystaceae | **Methylocystis** | - |
| | Proteobacteria | Alphaproteobacteria | Rhizobiales | Methylocystaceae | **Methylosinus** | - |
| | Proteobacteria | Alphaproteobacteria | Rhizobiales | Beijerinckiaceae | **Methylocapsa** | - |
| | Proteobacteria | Alphaproteobacteria | Rhizobiales | Beijerinckiaceae | **Methyloferula** | 1 |
| | Proteobacteria | Alphaproteobacteria | Rhizobiales | Beijerinckiaceae | **Methylocella** | 1 |
| | Proteobacteria | Gammaproteobacteria | **Methylococcales** | - | - | 21 |
| | Verrucomicrobiota | Methylacidiphilae | Methylacidiphilales | **Methylacidiphilaceae** | - | - |
| | NC10 | Methylomirabilacaea | Methylomirabilales | Methylomirabilaceae | **Ca. Methylomirabilis** | 1 |
| | Euryarchaeota | Methanomicrobia | Methanosarcinales | **ANME-2, ANME-3** | - | 2 |
| | Euryarchaeota | Methanomicrobia | Methanophagales | **ANME-1** | - | - |

This list is based on SILVA v128.
